# Control of Slc7a5 sensitivity by the voltage-sensing domain of Kv1 channels

**DOI:** 10.1101/2020.01.17.910059

**Authors:** Nazlee Sharmin, Shawn M. Lamothe, Victoria A. Baronas, Grace Silver, Yubin Hao, Harley T. Kurata

## Abstract

Many voltage-dependent ion channels are regulated by accessory proteins, although the underlying mechanisms and consequences are often poorly understood. We recently reported a novel function of the amino acid transporter Slc7a5 as a powerful regulator of Kv1.2 voltage-dependent activation. In this study, we report that Kv1.1 channels are also regulated by Slc7a5, albeit with different functional outcomes. In heterologous expression systems, Kv1.1 exhibits prominent current enhancement (‘disinhibition’) with holding potentials more negative than −120 mV. Disinhibition of Kv1.1 is strongly attenuated by shRNA knockdown of endogenous Slc7a5. We investigated a variety of chimeric combinations of Kv1.1 and Kv1.2, demonstrating that exchange of the voltage-sensing domain controls the sensitivity and response to Slc7a5. Overall, our study highlights additional Slc7a5-sensitive Kv1 subunits, and demonstrates that features of Slc7a5 sensitivity can be swapped by exchanging voltage-sensing domains.

**IMPACT STATEMENT:** The voltage-sensing mechanism of a subfamily of potassium channels can be powerfully modulated in unconventional ways, by poorly understood regulatory partners.

## INTRODUCTION

A wide array of ion channels underlie the ability of neurons to exhibit distinct and regulated patterns of firing (Gutman et al., 2005; Yu et al., 2005). Ion channel subtypes possess different voltage dependence, kinetics, sensitivity to signaling cascades, regulation by physiological ions, and other stimuli that contribute to the moment-to-moment and long term adaptability of electrical signaling in the nervous system. In contrast to the rich complexity of multi-protein complexes known to regulate many synaptic neurotransmitter receptors (Jacobi and von Engelhardt, 2018; Tomita, 2019), the vast majority of research on voltage-dependent potassium (Kv) channels has focused on mechanisms of voltage sensitivity. Studies of the Drosophila *Shaker* channel, the first cloned Kv channel, have generated a detailed understanding of core principles of voltage-dependent regulation (Bezanilla, 2008, 2006; Tempel et al., 1988; Timpe et al., 1988). In comparison, regulation of mammalian Kv channels by extrinsic factors such as accessory proteins or signaling cascades is less understood. It is noteworthy that the Kv channels are the most diverse ion channel gene family, with nearly 50 human genes known to encode pore-forming subunits, but there are a relatively small number of recognized and well-studied accessory proteins (Gutman et al., 2005).

Based on prior observations of variable Kv1.2 function in different cell types, we hypothesized that Kv1.2 is influenced by a variety of unidentified regulators (Abraham et al., 2019; Baronas et al., 2017, 2016, 2015; Rezazadeh et al., 2007). We have pursued the identification of novel regulatory proteins that may influence this widely used model Kv channel. We recently reported powerful effects of an amino acid transporter, Slc7a5, on the gating and expression of Kv1.2 (Baronas et al., 2018). Slc7a5 has been primarily studied in its role as a transporter of drugs and amino acids (Barollo et al., 2016; Dickens et al., 2017; Soares-da-Silva and Serrão, 2004), and also in the context of nutrient regulation of mTOR signaling (Nicklin et al., 2009; Wolfson et al., 2016), but is not recognized as a regulator of ion channel function. Recessively inherited Slc7a5 mutations were recently identified in patients with neurological symptoms including autism and motor delay. These traits were attributed to defective Slc7a5-mediated amino acid transport in endothelial cells in the blood brain barrier, where it is most prominently expressed (Kanai et al., 1998; Tărlungeanu et al., 2016). However, low levels of Slc7a5 have been reported in neurons, and other studies have suggested transport-independent functions of Slc7a5 in early development (Katada and Sakurai, 2019; Matsuo et al., 2000), indicating there may be additional unrecognized functions of Slc7a5.

Several structures of Slc7a5 have been recently reported, highlighting its conserved LeuT fold comprising 12 transmembrane helices (Lee et al., 2019; Yan et al., 2019), but there are few clues into the mechanisms underlying the powerful modulation of Kv1.2 by Slc7a5. Features of Slc7a5-dependent modulation of Kv1.2 currents include a pronounced shift (∼-50 mV) of the voltage-dependence of activation, along with significant potentiation after strong negative holding voltages, leading to channel ‘disinhibition’ that often reaches 10-fold enhancement of whole cell current. Strikingly, certain epilepsy-linked mutations of Kv1.2 are hypersensitive to Slc7a5-mediated modulation, leading to extraordinarily large (sometimes >100 mV) gating shifts (Baronas et al., 2018; Masnada et al., 2017; Syrbe et al., 2015). Although these effects are very prominent, the underlying structural determinants of the Slc7a5:channel interaction, its specificity among other Kv channels, and the role of Slc7a5 modulation in vivo, remain unclear.

In this study, we expanded our investigation of Slc7a5-mediated regulation to include another prominent neuronal potassium channel, Kv1.1. This channel subunit exhibits overlapping patterns of expression with Kv1.2 in the central nervous system, and often assembles with Kv1.2 into heteromeric channels (Coleman et al., 1999; Manganas and Trimmer, 2000; Shamotienko et al., 1997). Our findings demonstrate that Kv1.1 is especially sensitive to modulation by Slc7a5, such that even endogenous levels of Slc7a5 result in prominent effects on the channel. However, Kv1.1 channels exhibit different outcomes of Slc7a5 modulation relative to Kv1.2. We use this observation to probe how sequence differences between Kv1.2 and Kv1.1 govern Slc7a5 sensitivity, and identify the voltage-sensing domain as an important determinant.

## RESULTS

### Kv1.1 sensitivity to Slc7a5

We recently reported several powerful effects of Slc7a5 on the voltage-gated potassium channel Kv1.2 (Baronas et al., 2018). Among the prominent effects of Slc7a5 is a shift of the voltage-dependence of activation by roughly −50 mV (Fig. 1A, dashed lines, reproduced for comparison). We tested the effects of Slc7a5 co-expression with another prominent neuronal Kv1 channel, Kv1.1. Our initial characterization of Kv1.1 did not reveal a significant shift in the voltage-dependence of activation in the presence of Slc7a5 (Fig. 1A,B).

**Figure 1.**
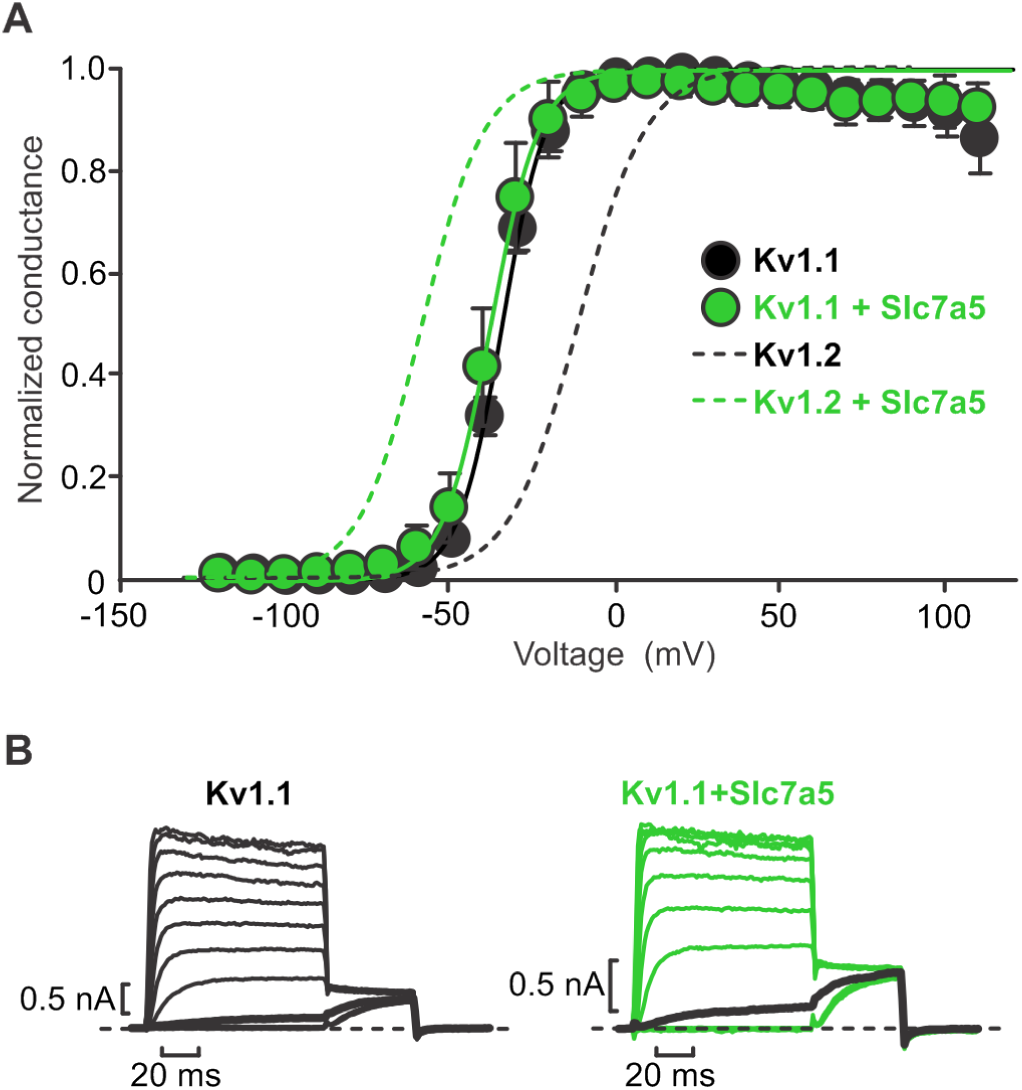
Slc7a5 has no effect on voltage-dependent activation of Kv1.1. (A) Conductance-voltage relationships were determined for indicated combinations of Kv1.1 and Slc7a5 expressed in LM mouse fibroblasts. Cells were stepped between −120 mV and +110 mV in 10 mV increments, and a tail current voltage of −20 mV. Dashed lines indicate previously reported conductance-voltage relationship in Kv1.2 ± Slc7a5 (Baronas et al., 2018). Fit parameter for Kv1.1 were (with +Slc7a5 in parentheses): V_1/2_ = −34.9 ± 0.3 mV (−37.5 +/- 0.2 mV); *k* = 6.9 ± 0.9 mV (7.3 ± 0.9 mV). (B) Exemplar records illustrating voltage-dependent activation of Kv1.1 ± Slc7a5. Current traces at −30 mV are bolded in black.

A second signature feature of Slc7a5 modulation of Kv1.2 was prominent disinhibition of current when membrane voltage was held at −120 mV or more negative. This behavior appears to be due to recovery from a non-conducting state, but the underlying mechanism is not yet fully understood. Exemplar records illustrating channel disinhibition are shown in Figures 2A and B, comprising intermittent 50 ms depolarizations to +10 mV from a holding potential of −120 mV (please note sweeps are concatenated for illustrative purposes − the interpulse interval was 2 s). Kv1.2 expressed alone exhibits no apparent disinhibition in response to this protocol (Fig. 2A, black) - disinhibition of Kv1.2 only becomes apparent when it is co-expressed with Slc7a5 (Fig. 2A, green,C,D). On average, we observed 5.1 ± 2.2 fold disinhibition of peak current for Kv1.2 co-expressed with Slc7a5, although in some cells as much as 11.5-fold disinhibition was observed (Fig. 2C,D). In contrast, we noticed prominent disinhibition for Kv1.1 (Fig. 2B), even in the absence of any co-transfected Slc7a5 (Fig. 2B,C). This effect was consistently observed, with an average disinhibition of 3.2 ± 1.2 fold. This effect, along with a suppression of Kv1.1 current density, became more prominent when Slc7a5 was co-expressed with Kv1.1, with a 6.1 ± 2.6 fold disinhibition (Fig. 2D).

**Figure 2.**
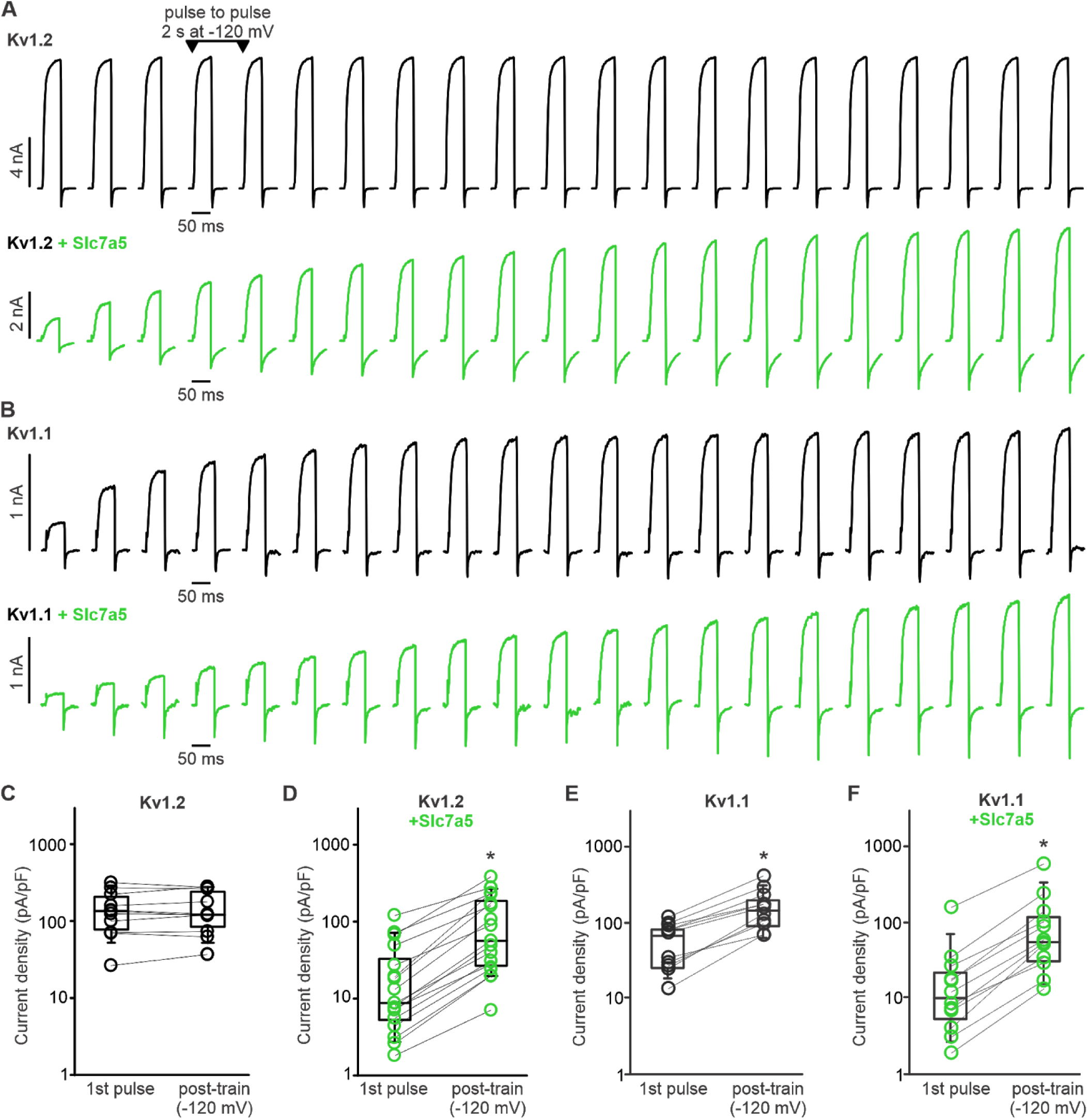
Kv1.1 exhibits prominent disinhibition in response to hyperpolarizing (−120 mV) voltage. (A,B) Disinhibtion of Kv1.2 (A) or Kv1.1 (B) was tested by delivering repetitive 50 ms depolarizations to +10 mV (every 2 s), with an interpulse holding voltage of −120 mV. When Kv1.2 is expressed alone, currents remain stable during this protocol (A), whereas Kv1.1 exhibits prominent disinhibition (B). (C-F) Cell-by-cell disinhibition measured before and after a hyperpolarizing pulse train to −120 mV is illustrated for indicated combinations of Kv1.1, Kv1.2 and Slc7a5. A prominent difference between Kv1.2 and Kv1.1 is that Kv1.1 exhibits disinhibition without a requirement for overexpression of Slc7a5 by co-transfection. Current density pre and post-train was compared using a paired t-test (* indicates p < 0.05). Kv1.2 (n=11, no statistical difference); Kv1.2 + Slc7a5 (n=16, p=0.001); Kv1.1 (n=11, p=0.004); Kv1.1 + Slc7a5 (n=13, p=0.003).

### Hypersensitivity of C-type inactivation

An additional powerful effect of Slc7a5 was a prominent acceleration of inactivation that was revealed in Kv1.2[V381T] (equivalent to *Shaker* position T449) mutant channels. Substitution to a Thr at this position influences susceptibility to C-type inactivation in *Shaker* channels (López-Barneo et al., 1993) and Kv1.2 (Goodchild et al., 2012). We tested inactivation of the analogous mutation Kv1.1[Y379T] (Fig. 3), and observed that Slc7a5 causes dramatic enhancement of the rate of inactivation (Fig. 3A). While the inactivation rate of Kv1.1[Y379T] was more pronounced in the presence of Slc7a5, it was also variable. We suspect that inconsistent expression of Slc7a5 in transfected cells was the origin of this variation, although there may be other regulatory factors in play that influence the response of Kv1.1 to Slc7a5. Overall, these findings demonstrate that Kv1.1 shares features of Slc7a5-mediated regulation with Kv1.2, although its response is not identical. Although we are not yet certain of the relationship between C-type inactivation and Slc7a5 modulation, this acceleration of inactivation is a prominent biophysical effect and highlights commonalities of Slc7a5 modulation of Kv1.1 and Kv1.2.

**Figure 3.**
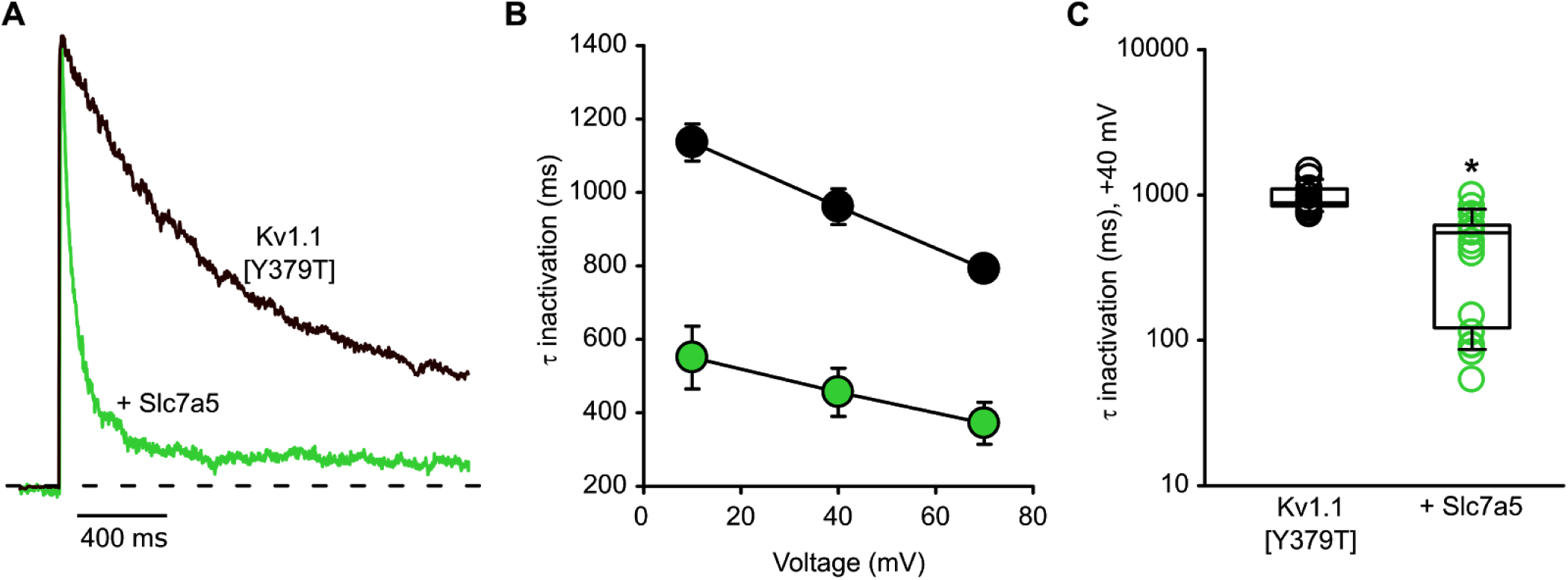
Pronounced inactivation of Kv1.1[Y379T] co-expressed with Slc7a5. (A) Kv1.1[Y379T] channels were expressed in mouse LM fibroblasts with Slc7a5, as indicated. Exemplar traces illustrate the time course of inactivation elicited by depolarization to +70 mV from a holding potential of −100 mV. Prior to this experiment, currents are disinhibited with a holding voltage of −120 mV, as described in Figure 2. (B) Mean time constant of inactivation for Kv1.1 ± Slc7a5, over a range of voltages. (C) Time constants of inactivation of Kv1.1 co-expressed with Slc7a5 (Kv1.1 τ = 960 ± 50 ms, n=17; Kv1.1 + Slc7a5 τ = 460 ± 60 ms, n=19). Inactivation time constants were compared with a student’s t-test, (* indicates p < 0.001).

### Knockdown and rescue of Slc7a5-mediated effects

Data thus far indicate that Slc7a5 modulation is distinct in Kv1.1 compared to Kv1.2, as there is no apparent shift in the voltage-dependence of Kv1.1 activation (Fig. 1). Also, Kv1.1 channels appear to be markedly more sensitive to Slc7a5 modulation, as they exhibit disinhibition consistent with Slc7a5-dependent modulation without a requirement for co-transfection of Slc7a5 cDNA. Therefore, Kv1.1 appears to be modulated by endogenous Slc7a5 expressed in LM cells. To test whether the disinhibition behavior of Kv1.1 was due to endogenous Slc7a5, we generated Slc7a5 knockdown cell lines (Slc7a5 ShR1, ShR2, ShR3 and ShR4) using lentiviral delivery and puromycin selection to maintain stable shRNA expression (Fig. 4 -supplement 1). Initial patch clamp experiments from these Slc7a5 shRNA cell lines exhibited significant cell-to-cell variability, but many cells had attenuated Kv1.1 disinhibition, along with larger currents relative to the parental LM cell line. This was especially apparent for ShR1 and ShR4 cell lines, suggestive of successful Slc7a5 knockdown (Fig. 4 -supplement 1). We further isolated individual clonal cell lines by serial dilution, from the ShR1 and ShR4 groups, and these clonal cell lines exhibited more consistent attenuation of Kv1.1 disinhibition (Fig. 4 -supplement 1,C). Most of the clonal cell lines also exhibited prominent reduction of Slc7a5 protein expression Fig. 4 -supplement 1,D). We selected a cell line (ShR4-1) with prominent knockdown of Slc7a5 confirmed by both real-time qPCR and Western blot (Figure 4A-C).

**Figure 4.**
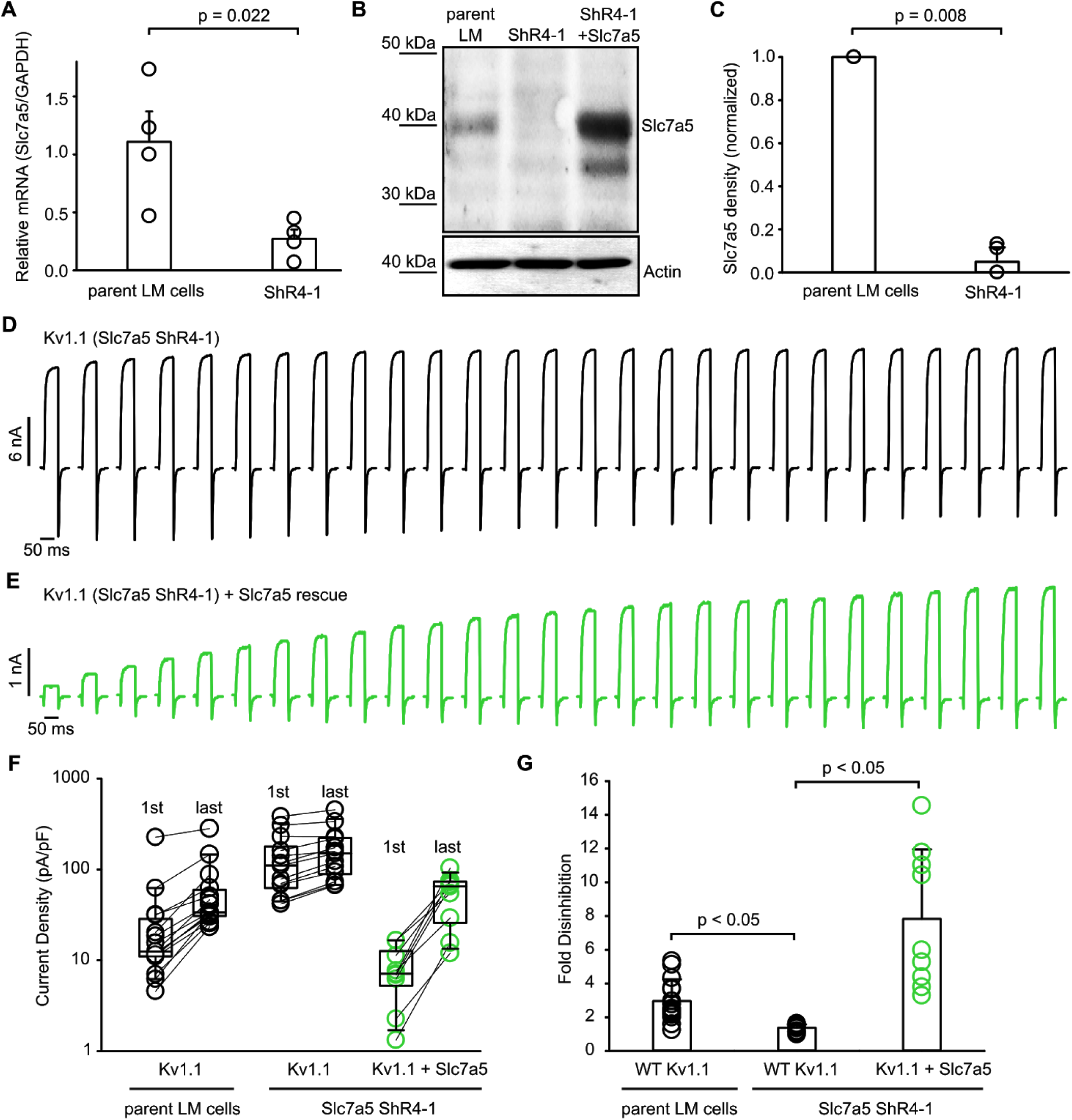
Modulation of Kv1.1 function by knockdown and rescue of Slc7a5. (A) Quantitative real-time PCR was conducted using RNA from parental mouse *LM* fibroblasts or a stable Slc7a5 shRNA knockdown mouse *LM* fibroblast cell line (ShR4-1) (n=4, statistical comparison with paired t-test). (B) Western blot of endogenous Slc7a5 in parental LM cells or Slc7a5 shRNA knockdown LM cells (ShR4-1). Actin was used as a loading control. (C) Densitometry measurements of Slc7a5 expression from parental and ShR4-1 LM cells (statistical comparison with paired t-test). Numbers in parentheses above the bars indicate the number of cells examined. (D,E) Exemplar current records illustrating disinhibition of Kv1.1 in response to a −120 mV hyperpolarizing voltage was measured as described in Figure 2, using the stable Slc7a5 shRNA knockdown (ShR4-1) mouse LM fibroblast cell line. In panel (E), Slc7a5 expression has been rescued by overexpression with a plasmid encoding human Slc7a5, which has 2 mismatches to the shRNA target sequence in the stable line. (F) Disinhibition from a −120 mV pulse train of Kv1.1 on a cell by cell basis in the parental LM cells or ShR4-1 cell line before or after Slc7a5 rescue (n = 9-15). (G) Bar graph depicting the fold disinhibition between the first and the last pulses of a −120 mV pulse train of Kv1.1 in parental LM cells (mean +/- S.D.; 2.96 ± 1.29) or ShR4-1 cells before (1.36 ± 0.20) or after (7.83 ± 4.12) Slc7a5 rescue (n = 9-15, Kruskal-Wallis multiple comparisons test, Dunn’s post-hoc test).

Expression of Kv1.1 in the ShR4-1 cell line resulted in a prominent increase in baseline current, and a lack of disinhibition relative to parental LM cells (Fig. 4D,F). An additional benefit of the ShR4-1 cell line is that the human Slc7a5 transcript is resistant to the shRNA sequence due to two base pair mismatches. Thus, we could rescue Slc7a5 expression in the ShR4-1 cell line using a human Slc7a5 cDNA (Fig. 4B,E-G), leading to the re-emergence of modulation observed in the parental LM cell line (Fig. 4E). Effects of exogenous expression of Slc7a5 in the ShR4-1 cell line included suppression of Kv1.1 current density, and far more prominent disinhibition (Fig. 4E-G). Overall, these findings indicate that Slc7a5 is essential for Kv1.1 disinhibition at negative voltages. Additionally, they highlight that Kv1.1 is particularly sensitive to Slc7a5 regulation, sufficient to observe modulation by endogenous levels of Slc7a5.

### Slc7a5 sensitivity is mediated by the voltage-sensing domain

Our findings illustrate that Kv1.1 and Kv1.2 can both be modulated by Slc7a5, but specific functional outcomes differ between these two channel types. While Kv1.2 appears to require higher expression of Slc7a5 to observe an effect (ie. it is less sensitive to modulation), it exhibits an identifiable ‘signature’ strong shift in the voltage-dependence of activation. In contrast, Kv1.1 appears to be sensitive to even endogenous levels of Slc7a5, but the gating effects are limited to current disinhibition (there is not a prominent gating shift).

We used these differences to investigate the determinants of Slc7a5 sensitivity using a chimeric approach. We swapped increasing segments of Kv1.2 with the corresponding sequence from Kv1.1, and measured the effects of Slc7a5 on current disinhibition and voltage-dependence of activation (Fig. 5). For each chimera, the effect of Slc7a5 on current magnitude before and after the disinhibition protocol is illustrated in Fig. 5A, and effects on voltage-dependent gating are presented in Fig. 5B-E. The N-terminus, S1, and S2 segments of Kv1.2 can be replaced while still preserving a large Slc7a5-mediated shift in voltage-dependent gating (Fig. 5A-D). However, further replacement of the S3 and S4 segments (and a small portion of the pore) led to a significant switch towards a Kv1.1-like phenotype (Fig. 5A,E). In terms of disinhibition, the Kv1.1S5/Kv1.2 chimera frequently exhibited prominent disinhibition of current in the absence of transfected Slc7a5 (Fig. 5A). In addition, the Slc7a5-mediated gating shift was largely attenuated in the Kv1.1S5/Kv1.2 chimera (Fig. 5E).

**Figure 5.**
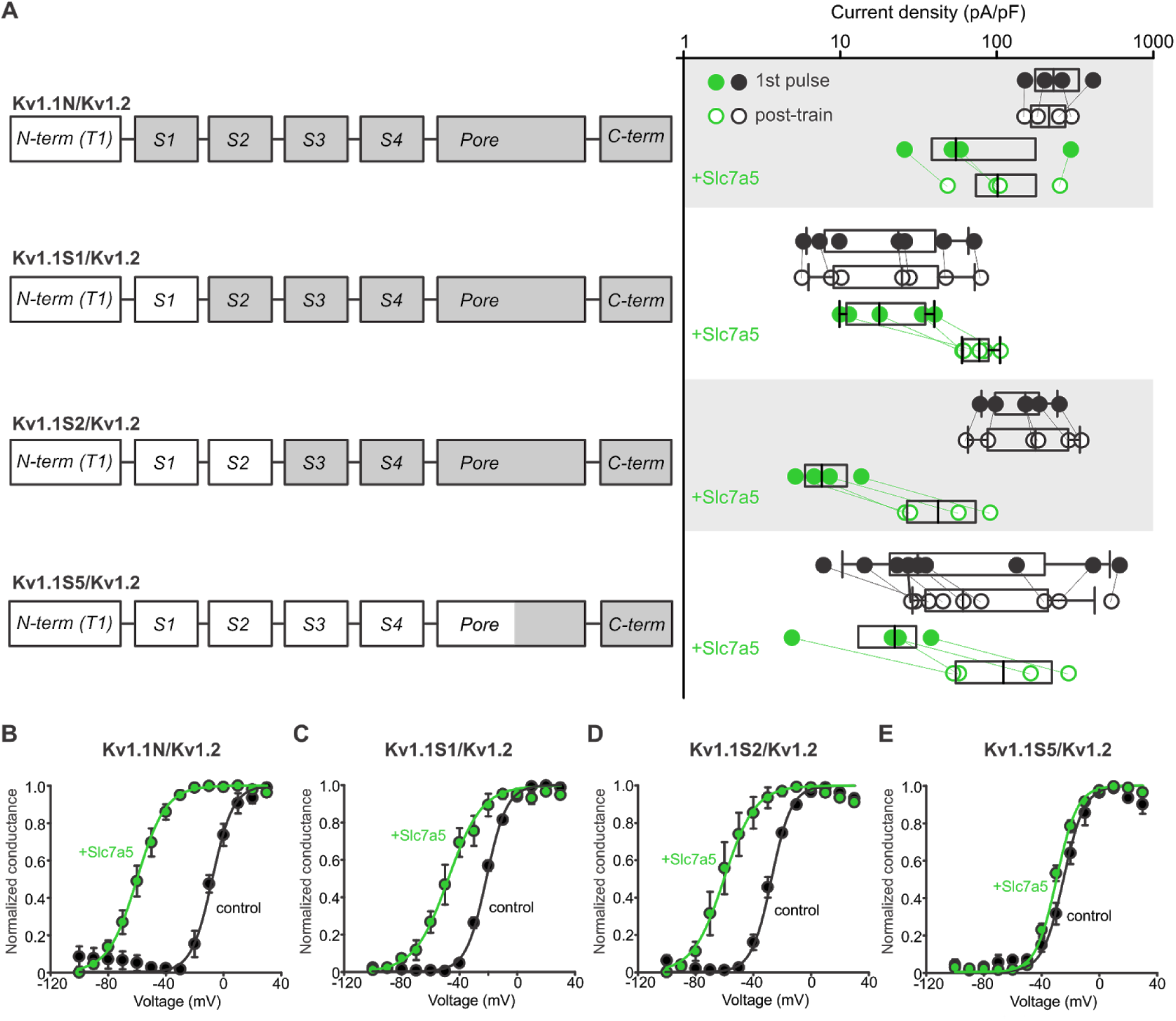
Chimeric analysis of Kv1.1 and Kv1.2 sensitivity to Slc7a5-mediated disinhibition and shifts of voltage-dependent activation. (A) Cartoons illustrate chimeric channel design, in which increasing segments of Kv1.1 (white) were introduced into Kv1.2 (grey), beginning with the N-terminus. Current disinhibition by a hyperpolarizing train to −120 mV was assessed as described in Figure 2, in the presence or absence of Slc7a5. (B-E) Conductance-voltage relationships were measured for all chimeric channels, in the presence and absence of Slc7a5. Gating parameters (+Slc7a5 in parentheses) for Kv1.1N/Kv1.2 were: V_1/2_ = −8.7 ± 2 mV (−61 ± 3 mV); *k* = 7 ± 1 mV (8.3 ± 0.5 mV); for Kv1.1S1/Kv1.2: V_1/2_ = −21.5 ± 0.7 mV (−42 ± 6 mV); *k* = 7.1 ± 0.2 mV (9 ± 1 mV), for Kv1.1S2/Kv1.2: V_1/2_ = −30.1 ± 0.4 mV (−53 ± 8 mV); *k* = 7.4 ± 0.2 mV (8.2 ± 0.4 mV), and for Kv1.1S5/Kv1.2: V_1/2_ = −25 ± 2 mV (−30 ± 2 mV); *k* = 7.4 ± 0.3 mV (7.2 ± 0.3 mV). Prominent shifts in voltage-dependent gating were observed in all chimeras except the Kv1.1S5/Kv1.2, comprising primarily the transmembrane domains of Kv1.1.

### VSD chimeras swap prominent features of Slc7a5 sensitivity

We further investigated the role of the voltage-sensing domain by swapping the S1-S4 segments of Kv1.1 and Kv1.2 (Fig. 6). Introduction of the voltage-sensing domain of Kv1.2 into Kv1.1 (Kv1.2VSD/Kv1.1) transferred all features of Slc7a5 modulation to Kv1.1. These chimeric channels exhibited a prominent Slc7a5-dependent shift in voltage-dependent gating, and current disinhibition, comparable to Kv1.2 (Fig. 6A,B). In addition, these features of Slc7a5 modulation were absent when the Kv1.2VSD/Kv1.1 chimera was expressed alone (Fig. 6A,B). The complimentary chimera that introduced the voltage-sensing domain of Kv1.1 into Kv1.2 (Kv1.1VSD/Kv1.2) did not have such clear cut effects, but certainly altered the Slc7a5 sensitivity of Kv1.2 (Fig. 6C,D). The Kv1.1VSD/Kv1.2 chimera exhibited prominent disinhibition of current even in the absence of Slc7a5, and a significantly attenuated shift in voltage dependence of activation when co-expressed with Slc7a5 (Fig. 6C,D).

**Figure 6.**
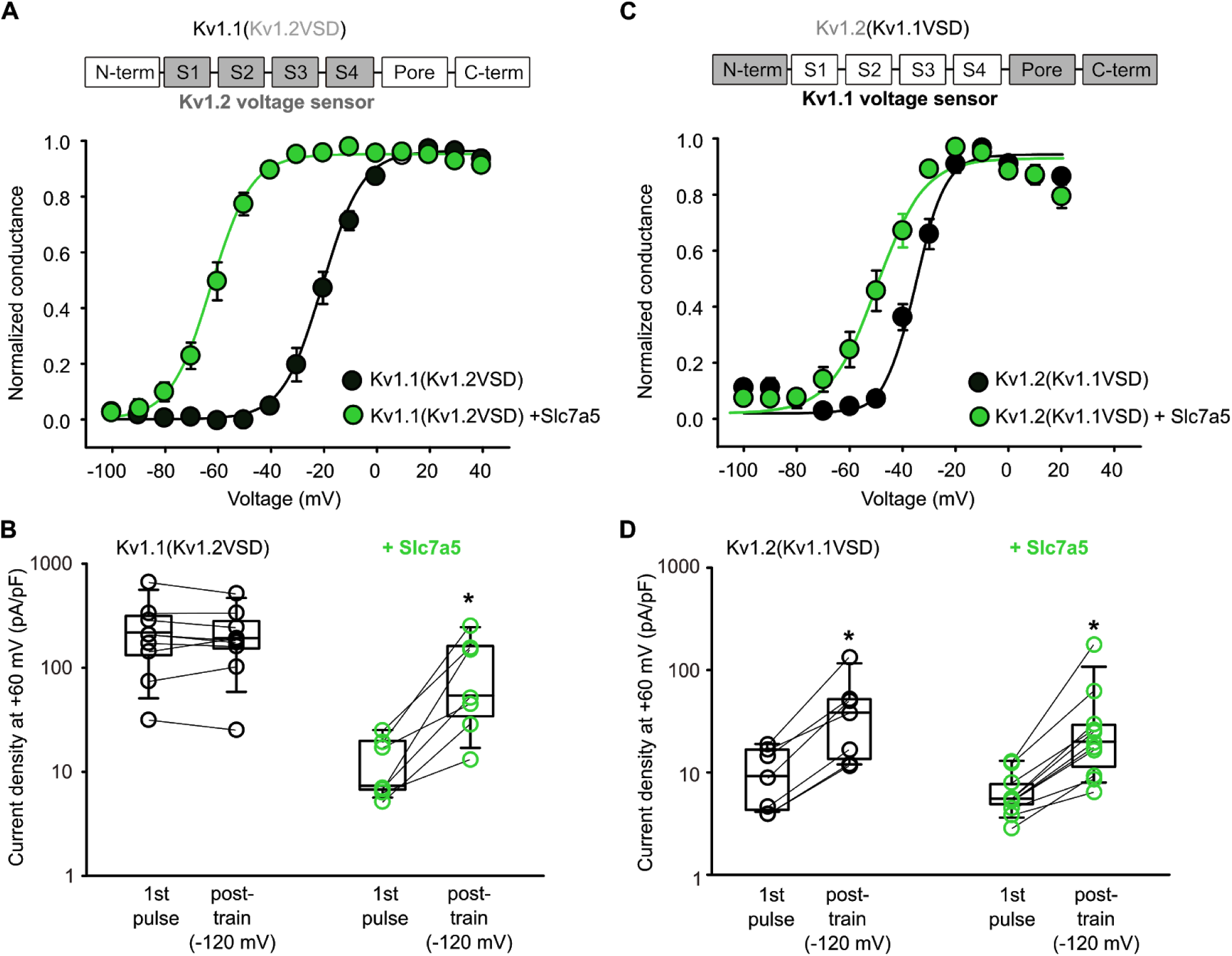
The voltage-sensing domain influences Slc7a5 sensitivity and response. (A,C) *Top:* Chimera design is shown with the voltage-sensing domains of Kv1.1(white) and Kv1.2 (grey) switched as indicated. Gating parameters for the resulting chimeras were (+Slc7a5 in parentheses), for Kv1.1(Kv1.2VSD): V_1/2_ = −19.7 ± 2 mV (−62 ± 2 mV); *k* = 7.5 ± 0.5 mV (7.7 ± 0.9 mV); for Kv1.2(Kv1.1VSD): V_1/2_ = −37 ± 2 mV (−53 ± 3 mV); *k* = 5.5 ± 0.4 mV (7.6 ± 0.9 mV). (B,D) Disinhibition of both chimeras was measured in the presence and absence of Slc7a5, in response to a −120 mV hyperpolarizing train, as described in Figure 2. Current densities pre- and post-train were compared using a paired t-test (* indicates p < 0.05). Fold disinhibition of Kv1.1(Kv1.2VSD) was 1 ± 0.1 (n=9, no statistical difference), and with Slc7a5 was 8 ± 3 (n=7, p=0.034). Fold disinhibition of Kv1.2(Kv1.1VSD) was 3.9 ± 0.6 (n=7, p=0.016), and with Slc7a5 was 4.3 ± 0.9 (n=11, p=0.001).

### Inhibition of Slc7a5 transport function or signaling does not prevent Kv1 modulation

Although we have previously demonstrated close proximity between heterologously expressed Kv1.2 and Slc7a5 via a BRET assay, it remains unclear whether Kv1.1 or Kv1.2 are regulated by a direct physical interaction with Slc7a5, or via some intermediary or a downstream signaling cascade. One possible prominent regulatory pathway linked to Slc7a5 is activation of mTOR via import of amino acids (Nicklin et al., 2009; Saxton et al., 2016; Wolfson et al., 2016), and so we considered whether a signal that relies on Slc7a5 transport activity underlies the prominent regulation of ion channel activity that we have observed. We tested whether pharmacological manipulation of Slc7a5 or mTOR activity would influence Slc7a5-mediated effects on Kv1.1, by incubating cells for 3-6 hours prior to recording, in either the commonly used Slc7a5 inhibitor BCH (2-amino-bicyclo[2,2,1]heptane-2-carboxylic acid), or the mTOR inhibitor rapamycin (Fig. 7). We observed that current disinhibition of Kv1.1 was insensitive to both of these treatments (Fig. 7A,B). We also collected conductance-voltage relationships to assess whether basal levels of Slc7a5 activity or a downstream signal could influence voltage-dependent activation of Kv1.1, however no effect of either BCH or rapamycin was observed (Fig. 7C,D). Although these experiments do not rule out involvement of an additional protein or signal for Slc7a5-dependent modulation of Kv1.1, our findings argue against the involvement of amino acid transport by Slc7a5, and downstream activation of mTOR.

**Figure 7.**
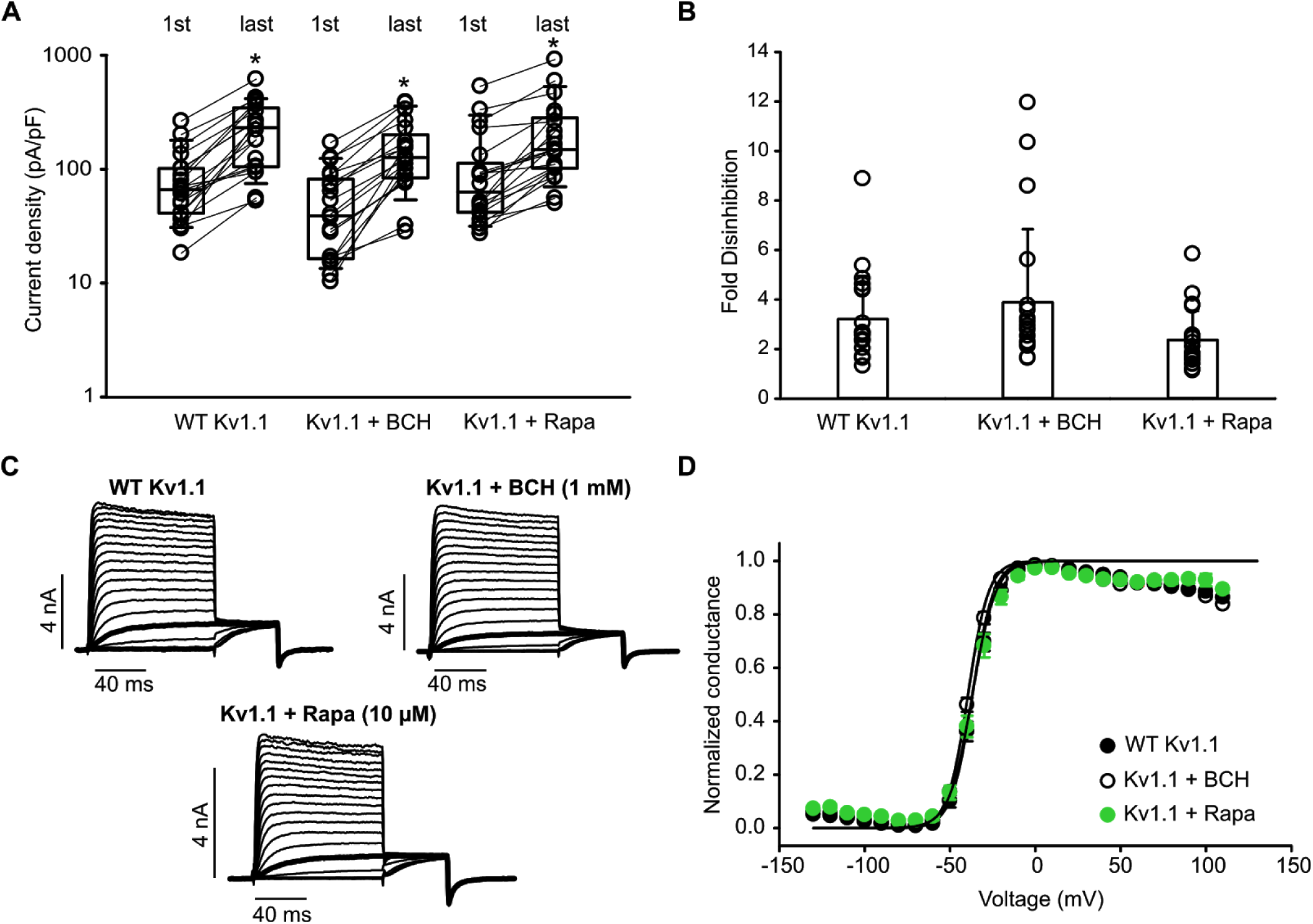
Slc7a5-mediated responses are not sensitive to inhibition of amino acid transport or mTOR signaling. **(A)** Cell-by-cell disinhibition measured before (1st pulse) and after (last pulse) a pulse train to −120 mV for 30s is illustrated for Kv1.1 in the absence and presence of 1 mM BCH, or 10 µM rapamycin. **(B)** Bar graph depicting the fold disinhibition between the first and the last pulses of a −120 mV pulse train of Kv1.1. WT Kv1.1: (mean ± S.D.; 3.2 ± 1.7); Kv1.1 + BCH (mean ± S.D.; 3.9 ± 2.9); Kv1.1 + Rapa (mean ± S.D.; 2.3 ± 1.2) (n = 19-20) **(C)** Representative current traces of the activation curves of Kv1.1 with the pulse to −30 mV highlighted. **(D)** Conductance-voltage relationship plots for Kv1.1 fit with a Boltzmann function in the absence and presence of 1 mM BCH, or 10 µM rapamycin. WT Kv1.1: V_1/2_ = −36.8 ± 1.2 mV; *k* = 5.6 ± 0.2 mV; n = 19; for Kv1.1 + BCH: V_1/2_ = −39.5 ± 0.7 mV; *k* = 5.7 ± 0.4 mV; n = 20; for Kv1.1 + Rapa: V_1/2_ = −36.9 ± 1.5 mV; *k* = 6.6 ± 0.5 mV; n = 20. Current densities pre- and post-train (panel A) were compared using a paired t-test (* indicates p < 0.05). Fold-disinhibition (panel B) in different conditions was compared using a one-way ANOVA, no statistical differences were detected.

## DISCUSSION

While the function of core alpha subunits of Kv channels has been investigated in depth, especially in the context of voltage-dependent gating, regulation of Kv channels by signaling pathways and regulatory proteins has been less widely studied. Our efforts to identify novel regulatory proteins of Kv channels led to the recent finding that Slc7a5, a widely studied amino acid transporter, exerts a powerful influence on gating and expression of Kv1.2 channels (Baronas et al., 2018). Prominent effects of Slc7a5 include suppression of channel expression, a −50 mV shift of voltage-dependent activation, powerful acceleration of inactivation of the Kv1.2[V381T] mutant, and marked disinhibition of current with negative holding potentials (indicating prominent but reversible suppression of Kv1.2 at physiological holding potentials). While powerful, the broader significance and underlying molecular mechanism(s) of these varied effects were not determined. In this study, we have continued to investigate Slc7a5 regulation of Kv channels and have begun to unravel some of these lingering questions.

We previously demonstrated some Kv1 subtype specificity of Slc7a5 regulation, as Kv1.2 was prominently affected by Slc7a5, while Kv1.5 was not (Baronas et al., 2018). Further investigation of other channel subtypes revealed that Kv1.1 exhibits many of the Slc7a5-mediated hallmark gating features reported for Kv1.2. Kv1.1 channels exhibit prominent disinhibition at negative voltages, along with pronounced inactivation of the Kv1.1[Y379T] mutant. Surprisingly, these features are prominent even in the absence of heterologous expression of Slc7a5 (Figs. 2). The influence of endogenous levels of Slc7a5 on Kv1.1 was confirmed using an shRNA knockdown approach, along with rescue by shRNA resistant Slc7a5 cDNA (Fig. 4).

We have taken some experimental steps to determine whether the Slc7a5-mediated effects on Kv1.1 are due to a direct interaction with the channel, or some indirect effect related to Slc7a5 function.

Slc7a5 is thought to influence mTOR activity, due to activation of the mTOR complex by amino acids like leucine (Nicklin et al., 2009; Saxton et al., 2016; Wolfson et al., 2016). Also, mTOR signaling has been shown to modulate Kv1.1 translation in dendrites, although this mechanism is unlikely to be involved here, as it was shown to rely on UTR elements that are not present in our Kv1.1 cDNA (Niere and Raab-Graham, 2017; Raab-Graham et al., 2006). We used a variety of pharmacological tools including direct inhibition of Slc7a5 transport (BCH), and mTOR inhibition (rapamycin), but found no effect on Kv1.1 regulation by Slc7a5. At present, we have not determined whether there is reciprocal modulation of Slc7a5 function by Kv channels. However, this possibility will continue to be explored, as certain combinations of Kv7 channels and myo-inositol transporters exhibit mutual regulation (Abbott et al., 2014; Manville et al., 2017; Neverisky and Abbott, 2017).

Auxiliary subunits can interact with voltage gated ion channels in a variety of ways. The canonical auxiliary subunits of the Kv1 channel family are the Kvβ subunits, which are soluble proteins that interact with a cytoplasmic scaffolding domain that is structurally distinct from the Kv1 transmembrane domains (Gulbis et al., 2000; Long et al., 2005). However, there is great diversity amongst known modulators of other Kv channels, including several transmembrane proteins. BK channels can be modulated simultaneously by both the BKβ and BKγ subunits, which are thought to associate with the pore-forming subunits via transmembrane domains (Gonzalez-Perez et al., 2015; Gonzalez-Perez and Lingle, 2019; Yan and Aldrich, 2012, 2010). Similarly, the widely studied KCNE subunits are transmembrane proteins that have been proposed to integrate into clefts between neighboring voltage-sensing domains to modulate channel function (Murray et al., 2016; Wang et al., 2012; Xu et al., 2013). Another Kv channel with prominent regulation of channel gating by multiple classes of regulatory proteins is the Kv4 family, which is sensitive to both KChIP (a soluble cytoplasmic protein) and DPP-like proteins (transmembrane), leading to modulation of channel expression and gating (Jerng et al., 2005; Kitazawa et al., 2015; Zagha et al., 2005). Although we have not yet collected direct evidence of an interaction between Slc7a5 and Kv1.1 or Kv1.2, our findings strongly suggest that Slc7a5 sensitivity is encoded by the voltage-sensing domains of these channels. In addition, our previous study demonstrated co-regulation of Kv1.2 by Slc7a5 and Kvβ subunits (Baronas et al., 2018), indicating the possibility that Kv1 channels might assemble with multiple accessory subunits simultaneously (similar to the putative Kv4 complex with KChIP and DPP-like proteins). It is also noteworthy that we have previously identified powerful regulation of Kv1.2 gating by extracellular redox conditions, by a mechanism that is not intrinsic to the Kv1.2 α-subunit (Baronas et al., 2017). This effect is vastly different from Slc7a5 modulation, indicating an additional regulatory mechanism, and a recent study has suggested a possible role for the sigma opioid receptor (Abraham et al., 2019). Thus, there are likely multiple unrecognized regulatory mechanisms that can strongly influence Kv1 channel gating, but it remains unclear how these pathways may interact in cell lines or native tissues.

In summary, our study highlights that Slc7a5 can modulate multiple targets in the Kv1 family, including Kv1.1 and Kv1.2. Moreover, we map Slc7a5 sensitivity to the voltage-sensing domain of Kv1 channels, suggesting that modulation involves interactions within the transmembrane segments, rather than cytoplasmic elements of these interacting proteins. We are hopeful that ongoing investigation of Kv channel regulatory proteins will broaden our understanding of voltage-gated channel function and regulation in vivo.

## MATERIALS AND METHODS

### Cell culture and expression

Mouse LM(tk-) fibroblast cells (ATCC), referred to throughout as LM cells, were used for patch clamp experiments, Western blots, and qPCR. Cells were maintained in culture in a 5% CO_2_ incubator at 37°C in DMEM supplemented with 10% FBS and 1% penicillin/streptomycin. Cells were split into 12-well plates to achieve 70% confluence the subsequent day, when they were transfected with cDNA using jetPRIME transfection reagent (Polyplus). Fluorescent proteins were used to identify transfected cells for electrophysiological recording. 6-10 hours after transfection, cells were split onto glass coverslips at a low density into 6-well plates, for electrophysiological recordings from single cells the following day. Electrophysiological recordings were done 24-36 hours after transfection.

Stable shRNA-mediated knockdown cell lines were generated by puromycin selection after lentiviral infection of parental LM cells. Puromycin resistant cells were plated in serial dilutions to isolate clonal cell lines. Knockdown cell lines were maintained in the same media as parental LM fibroblasts, along with 2.5 µg/mL puromycin to maintain expression of the shRNA.

### Potassium channel constructs

Kv1 channel cDNAs (human Kv1.1, rat Kv1.2, and various chimeric combinations of these channels) were expressed using the pcDNA3.1(-) vector (Invitrogen). Chimeric constructs of human Kv1.1 and rat Kv1.2 were generated using overlapping PCR approaches. N-terminal fragments of Kv1.1 were amplified by PCR using a 5’ flanking primer (Kv1.1Forward) and each of the following primers B-E. Additional channel segments from Kv1.2 were amplified using the reverse complement of primers B-E, and a 3’ flanking primer (Kv1.2Reverse).

Kv1.1Forward: GGGCTCGAGATGACGGTGATGTCTGGGGAG

B: GAAACTCATTTTCAGGCAGGGGGCGCTCCTCCTCCTTGATGAAGC

C: CTCTACGATGAAGAAAGGGTCTGTGAAGATGTTGGAATTGTAG

D: GCCACAATGTCAATGATGTTCATGATGTTTTTGAAGAAGTCCGTCTTGC

E: GGGGAACTGGGAATCTCGCTCATCCGCCTCGGCAAAGTACACTGCACTAG

Kv1.2Reverse: GGGGAAGCTTTTCAGACATCAGTTAACATTTTGG

Resulting full length channel fragments were then cloned into pcDNA3.1(-) using EcoRI and HindIII restriction digests and ligation. This lead to chimeric Kv1.1/Kv1.2 ion channels with break points at amino acid numbers, 147, 226, 256, 350 (Kv1.1 numbering, schematics of chimera design are shown in Fig. 5A). Constructs were all verified by diagnostic restriction digestion and Sanger sequencing (Genewiz, Inc., or University of Alberta Applied Genomics Core).

Chimeric switching of voltage-sensing domains was accomplished using similar overlapping PCR approaches, with break points corresponding to amino acid numbers 147 and 350 (Kv1.1) or 145 and 352 (Kv1.2). To generate the Kv1.2(Kv1.1VSD) chimera we used a previously made Kv1.1/Kv1.2 chimera (breakpoint at Kv1.1 amino acid 350, labeled Kv1.1S6/Kv1.2 in Fig. 6) and replaced the N terminus with sequence from Kv1.2 using the following primers:

Kv1.2 Forward: GGGCTCGAGATGACAGTGGCTACCGGAG

Kv1.2Nt-term Reverse: CCTTCTCGGGCAGAGGACGTTCTTCTTCCTTGATATAG

Similarly, the Kv1.1(Kv1.2VSD) chimera was made by switching the pore and C-terminus of our previously made Kv1.1N/Kv1.2 chimera (Fig 6) with corresponding Kv1.1 sequence using the following primers:

Kv1.1 Reverse: CCCGGATCCTTAAACATCGGTCAGTAGCTTG

Kv1.1 Pore Forward: AGAGAGCTAGGGCTGCTCATC

The Kv1.1[Y379T] mutation was generated with overlapping PCR approaches using the following mutagenic primers:

Kv1.1[Y379T]-Forward : CGGTCACATGACCCCTGTGACAATTG

Kv1.1[Y379T]-Reverse : CAATTGTCACAGGGGTCATGTCACCG

### Electrophysiology

Patch pipettes were manufactured from soda lime capillary glass (Fisher), using a Sutter P-97 puller (Sutter Instrument). When filled with standard recording solutions, pipettes had a tip resistance of 1-3 MΩ. Recordings were filtered at 5 kHz, sampled at 10 kHz, with manual capacitance compensation and series resistance compensation between 70-90%, and stored directly on a computer hard drive using Clampex 10 software (Molecular Devices). Bath solution had the following composition: 135 mM NaCl, 5 mM KCl, 1 mM CaCl_2_, 1 mM MgCl_2_, 10 mM HEPES, and was adjusted to pH 7.3 with NaOH. Pipette solution had the following composition: 135 mM KCl, 5 mM K-EGTA, 10 mM HEPES and was adjusted to pH 7.2 using KOH. Chemicals for electrophysiological solutions were purchased from Sigma-Aldrich or Fisher. BCH (Slc7a5 inhibitor 2-amino-bicyclo[2,2,1]heptane-2-carboxylic acid, Sigma-Aldrich) was stored as a 100 mM stock solution in 1N NaOH, and diluted to working concentrations each experimental day. Rapamycin (Alfa Aesar, Fisher Scientific) was stored as a 10 mM stock solution in DMSO and diluted to working concentrations each experimental day.

### Lentiviral vector construction and delivery

We generated shRNAs targeting four segments of mouse Slc7a5 (refseq: NM_011404), with the following sequences:

ShR1: GCAATATCACGCTGCTCAA

ShR2: GCAGAAGTTGTCCTTTGAA

ShR3: GGAACATTGTGTTGGCTTTG

ShR4: GCATTGGCTTCGCCATCAT

Negative control: GCAGTTATCTGGAAGATCAGG

Oligos were designed for hairpin formation and cloned into pLV-RNAi vector system (Biosettia, San Diego, USA. Cat#sort-B21) according to the manufacturer’s instructions. All constructs were confirmed by Sanger sequencing (Applied Genomics Core, University of Alberta). HEK293T cells were co-transfected with packaging vectors(Provided in the pLV-RNAi vector system) and an shRNA expression vector. Lentivirus was harvested by centrifugation of HEK293T cell culture supernatant 48 hours after transfection. LM fibroblasts cells were seeded in 6 well plate at about 30 % confluence the day before viral transduction. After 24 hours, media was replaced with fresh complete medium (DMEM with 10% FBS) containing 8 µg/ml polybrene and 0.5 mL of viral supernatant. After 24 hours of incubation, the virus-containing medium was replaced with fresh complete medium. After further incubation for 48 hours, puromycin (2.5 µg/mL) was added and maintained in culture for selection of transduced cells. For the generation of clonal shRNA-expressing cell lines, transduced cells were plated by serial dilution in 96 well plates, and wells with single cells were identified by visual inspection under a microscope. After expansion of individual clones, effectiveness of knockdown was assessed using Western blot and qPCR approaches.

### Western blot

Cell lysates from LM fibroblasts were harvested in NP-40 lysis buffer (1% NP-40, 150 mM NaCl, 50 mM Tris-HCl) with 1% protease inhibitor cocktail (Sigma, P8340), 3 days after transfection. Samples were separated on 8% SDS-PAGE gels, and transferred to nitrocellulose membranes using standard Western blot apparatus (Bio-rad). Running buffer composition was 190 mM Glycine, 25 mM Tris, 4 mM SDS. Transfer buffer composition was 20% methanol, 3.5 mM Na_2_CO_3_, 10 mM NaHCO_3_. Slc7a5 was detected using a rabbit polyclonal Slc7a5 antibody (1:500 dilution, KE026; Trans Genic Inc.) and HRP-conjugated goat anti-rabbit antibody (1:15 000 dilution, SH012; Applied Biological Materials). Chemiluminescence was detected using SuperSignal West Femto Max Sensitivity Substrate (Thermo Fisher Scientific) and a FluorChem SP gel imager (Alpha Innotech).

### Quantification of Slc7a5 mRNA expression using qPCR

Total cellular RNA was extracted from LM fibroblasts stably expressing shRNA constructs using an illustra RNAspin Mini kit (GE, UK). RNA concentration was assessed using a Nanodrop 2000c spectrophotometer (Thermo Scientific). Reverse transcription was performed using the SuperScript IV First-Strand Synthesis System (Invitrogen, USA), using Oligo(dT)20 primers to make cDNA. Real-time quantitative PCR was carried out with TaqMan® Fast Advance Master Mix (Applied Biosystems, USA) in an ABI 7900HT Fast Real Time PCR System (Applied Biosystems). Taqman probes used were: Slc7a5 (cat: Mm00441516_m1), and GAPDH as an internal control gene (cat: Mm99999915_g1), obtained from Thermo Fisher. The cycling protocol used an initial denaturation at 95 °C for 3 min; 40 cycles of denaturation at 95 °C for 3 s, annealing at 60 °C for 30 s. Data were analyzed using the 2^-ΔΔCT^ method (Livak and Schmittgen, 2001) and expressed as Slc7a5 normalized to GAPDH.

### Data analysis

Wherever possible, we have displayed data for all individual cells collected, in addition to reporting mean ± SEM or a box plot (where shown, box plots depict the median, 25^th^ and 75^th^ percentile (box), and 10^th^ and 90^th^ percentile (whiskers)). Conductance-voltage relationships were fit with a Boltzmann equation (Equation 1), where G is the normalized conductance, V is the applied voltage, V_1/2_ is the half activation voltage, and *k* is a fitted slope factor reflecting the steepness of the curve.

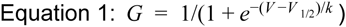

Conductance-voltage relationships were fit for each individual cell, and the extracted fit parameters were used for statistical calculations. Statistical tests are described in corresponding figure legends throughout the manuscript.

## ACKNOWLEDGEMENTS

This work was funded by a Canadian Institutes of Health Research Project Grant to HTK. SML was supported by a Rowland and Muriel Haryett Fellowship, University of Alberta Neuroscience and Mental Health Institute. VAB was supported by a Canadian Institutes of Health Research Vanier award. GS was supported by a Natural Sciences and Engineering Research Council USRA award. HTK was supported by a Canadian Institutes of Health Research Early Career Investigator award and salary support from the Alberta Diabetes Institute.

---

**Figure 4 - Supplement 1.**
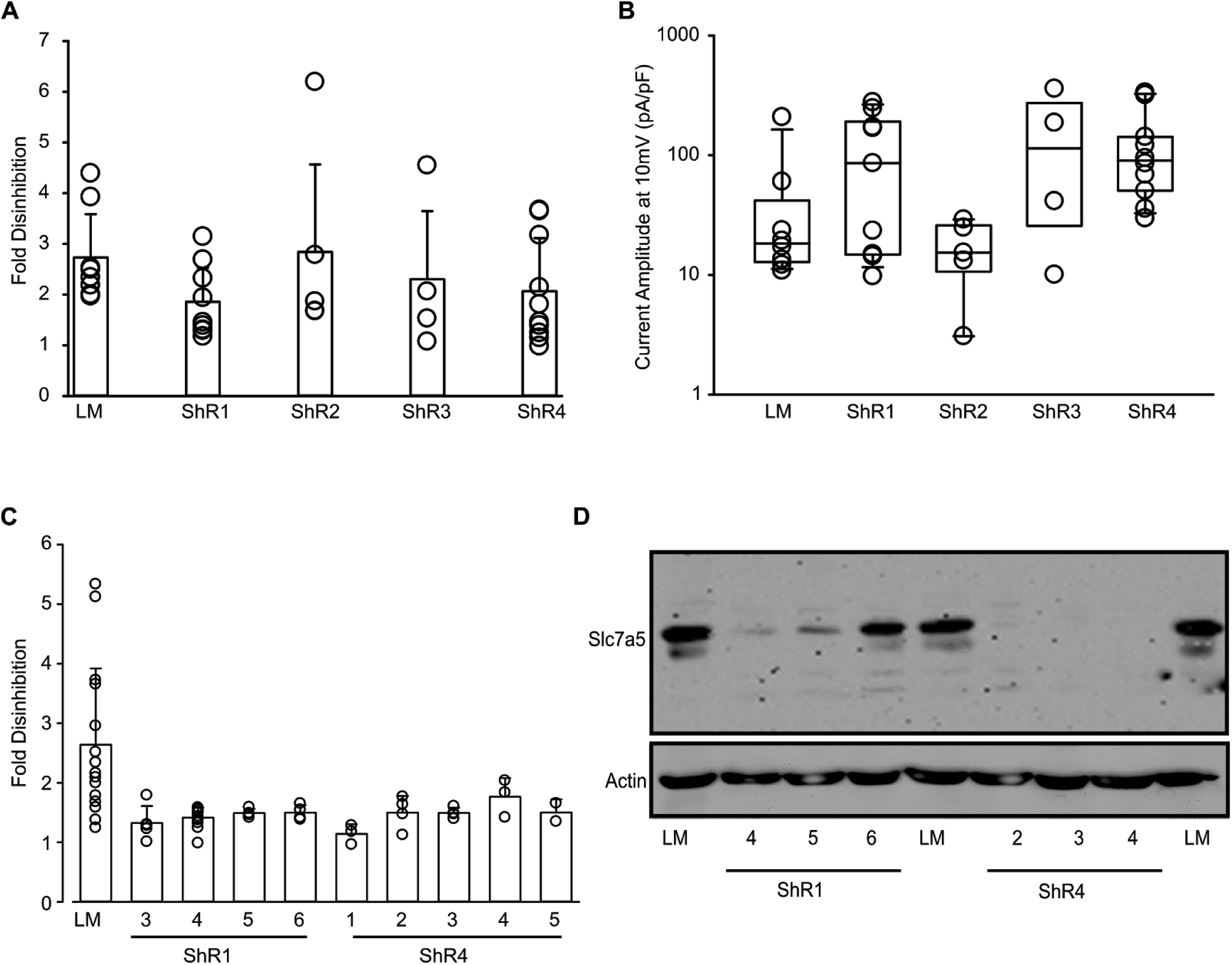
Generation of Slc7a5 knockdown cell lines. **(A,B)** Multiple cell lines expressing different shRNA constructs were transfected with Kv1.1 cDNA and tested for current disinhibition and current magnitude as described in Figure 2. **(C,D)** Multiple clonal cell lines were generated by serial dilution of the ShR1 and ShR4 cell lines, and screened for disinhibition of current (C) and Slc7a5 expression by Western blot (D). Further characterization of the ShR4-1 cell line is described in the main text.

